# Cre/LoxP-HBV plasmids generating recombinant covalently closed circular DNA genome upon transfection

**DOI:** 10.1101/2020.07.21.214171

**Authors:** Robert L. Kruse, Xavier Legras, Mercedes Barzi

**Affiliations:** Baylor College of Medicine, 1 Baylor Plaza, Houston, TX, USA 77030

**Keywords:** HBV, cccDNA, Cre/LoxP, drug discovery

## Abstract

New therapies against hepatitis B virus (HBV) require the elimination of covalently closed circular DNA (cccDNA), the episomal HBV genome. HBV plasmids containing an overlength 1.3-mer genome and bacterial backbone (pHBV1.3) are used in many different models, but do not replicate the unique features of cccDNA. Since the stable cccDNA pool is a barrier to HBV eradication in patients, we developed a recombinant circular HBV genome (rcccDNA) to mimic the cccDNA using Cre/LoxP technology. We validated four LoxP insertion sites into the HBV genome using hydrodynamic tail vein injection into murine liver, demonstrating high levels of HBV surface antigen (HBsAg) and HBV DNA expression with rcccDNA formation. HBsAg expression from rcccDNA was >30,000 ng/mL over 78 days, while HBsAg-expression from pHBV1.3 plasmid DNA declined from 2,753 ng/mL to 131 ng/mL over that time in immunodeficient mice (P<0.001), reflective of plasmid DNA silencing. We then cloned Cre-recombinase in cis on the LoxP-HBV plasmids, achieving plasmid stability in bacteria with intron insertion into Cre and demonstrating rcccDNA formation after transfection *in vitro* and *in vivo*. These cis-Cre/LoxP-HBV plasmids were then used to create HBx-mutant and GFP reporter plasmids to further probe cccDNA biology and antiviral strategies against cccDNA. Overall, we believe these auto-generating rcccDNA plasmids will be of great value to model cccDNA for testing new therapies against HBV infection.

## Introduction

Hepatitis B virus (HBV) chronically infects over 257-270 million patients worldwide (2015 World Health Organization estimate) [1], causing significant morbidity and death, including a substantial burden of 800,000 – 1.4 million people in the United States [2]. There is currently no cure for HBV infection; a key barrier is thought to be the stability of covalently closed circular DNA (cccDNA), the HBV genome, inside host hepatocytes under current therapies [3]. Indeed, the half-life of HBV cccDNA has been estimated from months to years [4]. Therapies to cure HBV would ideally trigger silencing or degradation of cccDNA.

Unfortunately, cccDNA does not form in mice, the most common animal model for biomedical research [5]. HBV cccDNA naturally forms in humans and chimpanzees during infection, and in other species (cynomolgus macaque, rhesus macaque, and pigs) after introduction of the sodium taurocholate cotransporting polypeptide (NTCP) receptor [6]. As an alternative, researchers have developed recombinant cccDNA (rcccDNA), using recombinases to excise the bacterial backbone. The recombination sequence (ex: LoxP) has been inserted into the HBV genome by various strategies, including identification tolerated insertion sites of the recombination sequence upstream of the Core and PreS1 proteins [7,8], or within an artificial intron introduced the HBV surface antigen (HBsAg) gene [9-11]. These rcccDNA templates have been prepared by the co-transfection of Cre recombinase into target cells [9], introducing Cre into cells harboring integrated LoxP-HBV [10], or by introduction into mice expressing Cre recombinase [9,11]. Alternatively, rcccDNA can be made within *E. coli* themselves, like current minicircle DNA molecules [7,8]. Regardless of strategy, these artificial cccDNA molecules have shown similar histone organization and DNA supercoiling to native cccDNA, and superior stability, persistence, and expression profile compared to HBV plasmids in mouse models [7-11].

While providing the field with tools that mimic HBV cccDNA, these strategies still possess several limitations. Recombinant cccDNA containing introns create significantly larger than wildtype size genomes. Moreover, introns could potentially increase HBsAg expression above wildtype HBsAg levels [12]. Relying on co-transfection of Cre and LoxP-HBV into cells or animals creates a mixture of excised and unexcised genomes, which can complicate experimental readout. This disadvantage is not seen in studies generating rcccDNA inside *E. coli* [7,8], but minicircle production requires special strains and culture complexity not widely used in labs [13]. Thus, there is a compelling need to generate cccDNA in a controlled and reproducible manner within every cell, while also allowing for ease of modification sites of the HBV genome in order to control to study HBV biology.

Herein, we describe a system of creating rcccDNA using LoxP sites inserted into multiple tolerated sites in the genome, resulting in HBV genome sizes similar to wildtype virus and serving as mutual controls for disruption of viral function in experiments. We also describe the construction of a single plasmid harboring Cre and LoxP-HBV, affording rcccDNA formation in cells or in mice, in an easy-to-use plasmid vector that can be grown in routine *E. coli* cultures and purified by established processes.

## Materials and Methods

### Plasmid Construction

#### Insertion Strategy

The HBV genome was analyzed for sites that could tolerate insertion of the LoxP sequence into the genome. One region identified for insertion was in the overlapping PreS and spacer domains of the hepatitis B surface antigen (HBsAg) and polymerase (Pol), respectively. The spacer domain has been previously shown to tolerate a wide variety of deletions and insertions while maintaining polymerase activity [14,15]. The key PreS domain amino acid residues for entry have also been finely mapped [16]. Furthermore, mutations that abolish entry motifs in the PreS domain may be less important since these plasmids are artificially introduced into human liver cell lines and mice, respectively, and a spreading, entry-based infection is not seen in either system. A schematic of the LoxP insertion into the PreS1 and PreS2 domains is shown in **Fig. 1A**, with a more detailed schematic of their relation to HBV promoters before excision provided in **Fig. S1**. Note that for the LoxP insertions into these sequences, both the surface and polymerase open reading frames (ORFs) must be maintained to provide for replication and virion secretion, which necessitated the use of a modified LoxP sequence [17]. For the PreS1 construct, the crucial region at the n-terminus of the PreS1 domain [18] engaging the NTCP entry receptor was replaced by LoxP recombination sequence, meaning the virus made by the recombinant cccDNA molecules is non-infectious.

**Figure 1.**
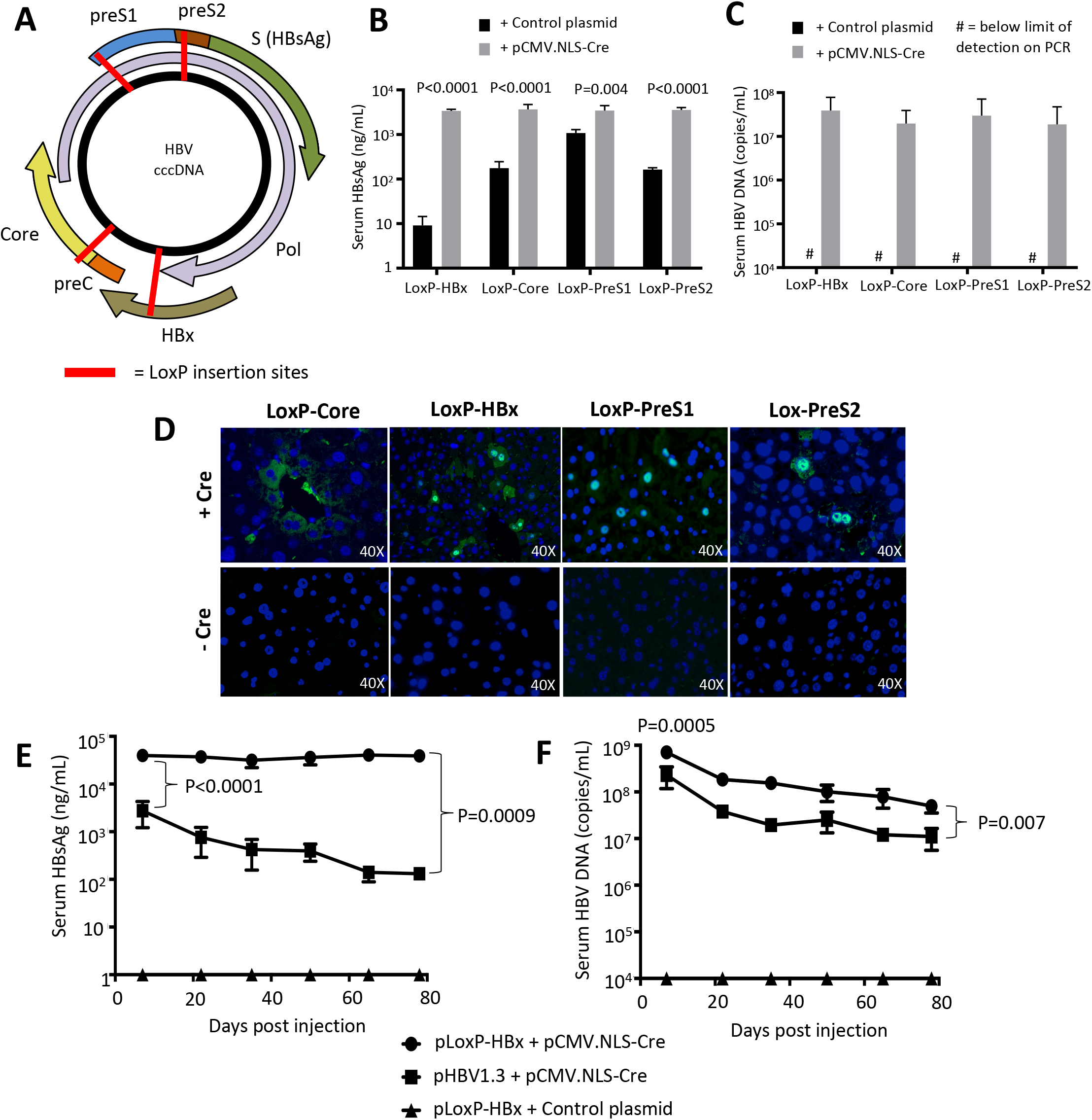
Recombinant cccDNA can be formed with LoxP insertion points resulting in similar production of viral biomarkers and is stable in mice long-term compared to HBV plasmids. **(A)** Four different LoxP insertion sites were identified in the HBV genome and predicted to not interfere with important viral functions. 15 μg of pLoxP-HBV plasmid (HBx, Core, PreS2, PreS1) were injected with 5 μg pCMV.NLS-Cre or Control plasmid into female NSG mice. HBsAg levels **(B)** and serum DNA levels **(C)** were measured at one week post-injection, showing increased expression with presence of Cre (n=4). **(D)** Immunofluorescence staining for Core expression in the same mice revealed Core staining only with the presence of Cre recombinase. **(E**,**F)** Long-term expression of rcccDNA generated from the LoxP-HBx construct was assessed in male NSG mice. 15 μg LoxP-HBx (n=4) or pHBV1.3 (n=3) were hydrodynamically injected with 5 μg pCMV-NLS-Cre or control plasmid. Serum HBsAg levels **(E)** and serum DNA levels **(F)** were monitored with elevated HBV expression seen with rcccDNA. Data represent mean ± standard error of mean (SEM); statistical significance (P<0.05).

Two other tolerant regions were designed in the hepatitis B core (HBc) protein and hepatitis B X protein (HBx), respectively (**Fig. 1A, S1**). It is known that HBV genotype G virus has a 12 amino acid insertion near the n-terminus of the HBc protein [19]. This insertion has been shown to be a peptide protruding out from the capsid surface, otherwise not interfering with capsid assembly [20]. The insertion region in the HBx protein was designed between the two motifs necessary for transcriptional trans-activation [21], but after the polymerase coding sequence ends and before the beginning of the enhancer II region, part of the core promoter [22]. Amino acid sequences including the LoxP coding sequence inserted into the designated HBV proteins is given in **Fig. S2C**.

#### Construction

The plasmid pSP65-ayw1.3 (named pHBV1.3 in this study) encoding a genotype D, serotype ayw HBV genome (gift of Dr. Stefan Wieland, University of Basel, Switzerland) was used as a template to make LoxP-HBV DNA plasmids. Gene synthesis (BioBasic, Amherst, NY) was employed to make a modified cloning site and ribozyme sequence, SmaI – nucleotide spacer - SacII – hammerhead ribozyme - HDV ribozyme – NheI,PacI (see **Fig. S2B**) for ribozyme sequence information), and cloned into pUC57 between EcoRI and HindIII forming pUC-Rbz. To create the LoxP-HBV cassettes, a strategy of triple ligation was performed into the pUC-Rbz. Two fragments of HBV genome would be generated by PCR, the external primers adding LoxP sites and the internal primers using a restriction site located within the HBV. General scheme of triple ligation to form pLoxP-HBV : 5’ Smal – LoxP – HBV sequence - Internal HBV restriction site + 5’ Internal HBV restriction site – HBV sequence – LoxP – SacII 3’ into 5’ SacII – Ribozymes – pUC57 – SmaI - 3’. The internal restriction site for HBc and HBx insertions was XhoI, and the internal restriction site for PreS1 and PreS2 insertions was NcoI. See **Table S1** for LoxP-HBV primer listing.

The subsequent pLoxP-HBV plasmids were named by the insertion point into the HBV genome: pLoxP-Core, pLoxP-PreS1, pLoxP-PreS1, pLoxP-HBx. Sequence information for the location of the LoxP insertion in the HBV genome at the DNA and amino acid levels is provided in **Fig. S3** and **Fig. S2C**, respectively.

To create the Cre/LoxP-HBV series of plasmids, a nuclear localization sequence (NLS)-Cre recombinase gene was cloned with an internal intron in order to prevent Cre expression in *E. coli* during plasmid propagation. The pCMV.NLS-Cre plasmid, derived from pcDNA3, contained an SV40 NLS appended to the n-terminus of Cre recombinase and was used as the starting material. A canonical intron site sequence (CAG/G) located in the Cre recombinase gene between BamHI and EcoRV sites in the Cre open reading frame was identified. A DNA fragment was synthesized (IDTDNA, Coralville, Iowa) that would introduce the human growth hormone intron in the Cre coding sequence, and cloned into this vector between BamHI and EcoRV (portion of sequence given in **Fig. S4A**,**B**). The pCMV.NLS-Cre(intron) plasmid, was then digested with ScaI to EcoRI (blunted) and the CMV-NLS(intron) fragment was cloned into pLoxP-HBV vectors cut with ScaI and SmaI, yielding pCre/LoxP-HBV vectors, or pCre/LoxP-HBV (with similar notation as above for the individual insertion points). The inserted fragment would be expected to use the internal polyadenylation sequence in the HBV genome in order to produce transcripts.

We obtained pHBV1.2, pHBV1.2*7, and pSI-X (gift of Betty Slagle, Baylor College of Medicine) that were previously described [23], with the latter two representing HBV genome with ablated HBx protein production and pSI-X expressing HBx under SV40 late promoter. In order to make a recombinant cccDNA molecule with no expression of HBx protein, pHBV1.2*7 plasmid was digested with NcoI to PsiI sites, and ligated into the same site in the pCre/LoxP-PreS2, forming pCre/LoxP-PreS2*7. The plasmid, pCMV-Gaussia (ThermoFisher, Waltham, MA), was used an empty control plasmid for the experiments.

The plasmid pCre/LoxP-PreS2 was used to make cccDNA reporter plasmids using green fluorescent protein (GFP). In the design, pCre/LoxP-PreS2 has GFP placed over the small HBsAg gene interrupting polymerase ORF, while also using the PreS2/S promoter to drive GFP expression. Primers amplified GFP adding EcoRI and SpeI sites, and cloned in between those same sites in pCre/LoxP-PreS2. The resultant plasmid was named pCre/LoxP-HBV-GFP(S). See **Table S1** for GFP primers utilized.

All plasmid cloning was verified by sequencing (Lone Star Labs, Houston, Texas). Cloning schemes and alignment were determined using Serial Cloner 2-6-1. To check for plasmid size where indicated, restriction digestion using restriction enzymes (New England BioLabs, Ipswich, MA) run on a 1% agarose gel was employed.

#### Cell culture assays

To test for the efficacy of the pCMV-NLS-Cre(intron) construct in successful intron splicing and Cre recombinase function, a transfection assay was performed with a pCAG-LoxP-STOP-LoxP-ZsGreen [24] (gift from Pawel Pelczar; Addgene plasmid #51269) that would serve as a reporter for Cre recombination. A pcDNA3-GFP plasmid served as the transfection control. Equal amounts of plasmid (2 μg total) were co-transfected with 3 μL 25kDa PEI (VWR, Radnor, PA) into 293T cells into a 24-well plate.

To test for excision and the fidelity of recombination, the pCre/LoxP-HBV plasmids (4 μg) were transfected with 6 μL PEI into 293T cells in 6-well plate. After 48 hours, cells were harvested. Pellets were collected from the cells, and circular DNA isolated using the ZR Plasmid Miniprep kit (Zymogen, Irvine, CA). PCR reactions over the recombination junction were set up using isolated DNA in order to assess for fidelity of Cre recombination. See **Table S1** for sequence of rcccDNA primers utilized.

To test for GFP reporter knockdown, a transfection assay was performed pCre/LoxP-HBV-GFP(S) to test for GFP knockdown with CRISPR/Cas9. The plasmid, pX330-U6-Chimeric_BB-CBh-hSpCas9 [25] (gift from Feng Zhang; Addgene plasmid # 42230), was cloned with a guide RNA 21 to form pCas9-gRNA21 [26]. The pCMV-Gaussia Luc plasmid served as the control. Equal amounts of plasmid (2 μg total) were co-transfected into 293T as described above.

Fluorescence was visualized using the Leica Microsystems (Wetzlar, Germany) fluorescent microscope and software program.

#### Animal experiments

Animal experiments followed a protocol approved by the Baylor College of Medicine Institutional Animal Care and Use Committee. Hydrodynamic tail vein injection was used to transfect plasmid DNA in murine livers according to established protocols [27]. For hydrodynamic tail vein experiments, NOD scid gamma (NSG), NOD-*scid* IL2Rg^null^, mice (Jackson Laboratory, Bar Harbor, ME) were utilized unless otherwise indicated in order to avoid immune responses against the HBV transgene that might have developed during long-term analysis. Testing for pLoxP-HBV functional screening utilized an pCMV.NLS-Cre plasmid from our lab to test for excision.

#### Mouse blood and tissue collection

For all blood draws, retro-orbital bleeding was used to obtain blood, after which blood was spun down for 30 minutes at 2.3G and serum collected. DNA was extracted from tissues using the DNeasy Blood & Tissue kit (Qiagen) and tested for the rcccDNA Cre excision products. Primers were also designed (see **Table S1** for sequences). PCR was performed according to manufacturer’s protocol (Phusion High-Fidelity PCR kit, New England Biolabs, Ipswich, MA).

#### HBV analysis of blood and tissue

Immunofluorescent staining for HBV core protein and determination of serum HBsAg and HBV DNA levels were performed as described previously [28,29].

#### Statistical analysis

Statistical analysis was performed using GraphPad Prism 7 software (GraphPad Software, Inc., La Jolla, CA). Data measurements are presented as mean ± standard error of mean (SEM). Mean differences were tested using appropriate tests including unpaired, parametric, two-tailed t-tests. Significance level used was P<0.05, unless otherwise specified.

## Results

### Identifying and constructing LoxP sequence insertion sites in the HBV genome

Through careful analysis of the HBV genome, we identified multiple sites (PreS1, PreS2, HBx, Core) in the HBV genome that may be tolerant to nucleotide insertions containing the LoxP sequence into open reading frames (ORFs) of the virus (**Fig. 1A**). The genome site selection is complicated by multiple ORFs and RNA/DNA regulatory elements throughout the HBV genome. We chose a nucleotide insertion strategy in order to avoid deleting important protein domains essential for the function of HBV proteins, strategizing that the extra 12 amino acids encoded by the LoxP site would just form an extra loop or linker in a protein and not cause interference. Using PCR to amplify segments of the HBV genome and primers adding LoxP sites to the ends of the fragments, the four insertion variants were constructed (**Fig. S1**) to form LoxP-HBV genomes (**Fig. S2A**). Hepatitis delta virus (HDV) ribozyme [30] and hammerhead ribozyme [31] were added at the 3’ end of all the LoxP-HBV genomes (**Fig. S2B**) to effectively cut all HBV transcripts resulting in degradation and lower protein production in the linear LoxP-HBV form (**Fig. S2A**). Complete protein and DNA sequences of the insertion sites are provided in **Fig. S2C, S3**. Additional information on the rationale for each insertion site selection is provided in the **Supplementary Methods**.

### HBV genomes with LoxP sequences in different sites successfully recombine and express viral antigens

We first tested if the different LoxP-HBV variants would successfully recombine after co-transfection with Cre recombinase to form rcccDNA, allowing for expression of HBV virions (measured by serum DNA levels) and HBsAg protein. When pLox-HBV plasmids were co-injected with Cre recombinase, we found that serum DNA (range: 1.87×10^7^ – 3.89×10^7^ copies/mL) could only be measured when pCMV-NLS-Cre was provided, and peak HBsAg levels (range: 3,380 ± 148 ng/mL – 3,652 ± 473 ng/mL) were only observed when Cre was provided as well (P<0.005) (**Fig. 1B**,**C**). HBsAg expression was also detected in the linear forms of the PreS1 (1,078 ± 105 ng/mL), PreS2 (163 ± 7 ng/mL), and Core (175.5 ± 31 ng/mL) LoxP-HBV insertions, since the HBsAg sequence was unperturbed in these vectors and some mRNA was likely expressed even with 3’ end ribozyme cleavage. The LoxP-HBx plasmid had the lowest basal HBsAg expression (9 ± 2 ng/mL) consistent with its critical importance in activating global HBV expression [32]. Immunostaining for Core antigen in the liver was only positive in mice which were co-transfected with the Cre recombinase (**Fig. 1D**). The pLoxP-Core plasmid had a more cytoplasmic Core immunostain, which may be due to the presence of the 12 amino acid insertion on the surface of the Core protein.

### Recombinant cccDNA yields higher and longer-term expression than HBV plasmids in mice

We next compared the expression of rcccDNA versus HBV overlength 1.3-mer plasmid (pHBV1.3) after hydrodynamic tail vein injection in immunodeficient mice. For this purpose, pLoxP-HBx plasmid was chosen, given its minimal background HBsAg expression. We injected pLoxP-HBx plasmid with and without Cre, along with pHBV1.3 as a control. pHBV1.3 demonstrated an initial peak of expression and then a prolonged decay of expression (**Fig. 1E**). rcccDNA (pLoxP-HBx + Cre) exhibited an initial 14-fold higher expression of HBsAg compared to the pHBV1.3 at one week (plasmid, 2,753 ± 890 ng/mL vs. rcccDNA, 39,763 ± 690 ng/mL, P<0.0001). HBsAg levels in rcccDNA were maintained for 78 days (38,953 ± 4,691 ng/mL at day 78), while pHBV1.3 saw its HBsAg level decline to 131 ± 20 ng/mL during this period (**Fig. 1E**), resulting in a 297-fold difference between rcccDNA and pHBV1.3 at day 78 (P=0.0009). Similarly, rcccDNA also yielded 3-fold, significantly higher serum DNA levels than pHBV1.3 at one week post-injection (7.16×10^8^ vs 2.32×10^8^ copies/mL, P=0.0005) (**Fig. 1F**). This difference was stable during the study period, with a 4.5-fold, significantly higher serum DNA levels of rcccDNA than pHBV1.3 at day 78 as well (5.00×10^7^ vs. 1.11×10^7^ copies/mL, P=0.007).

### Construction of a LoxP-HBV plasmid with Cre recombinase in cis

To make the system more robust, we cloned a Cre recombinase expression cassette and each LoxP-HBV sequence onto the same plasmid assuring that transfected LoxP-HBV plasmids always have Cre expression in the same cell to convert into rcccDNA form. In order to avoid premature excision and loss of LoxP cassettes during propagation in bacteria due to basal low-level expression of Cre in *E. coli* [33], we inserted a mammalian intron into the Cre ORF on a *cis*-plasmid with the LoxP-HBV genome. This intron can be successfully spliced in mammalian cells, but not in bacteria, thereby elimination Cre expression in *E. coli*.

The human growth hormone (hGH) intron was inserted into an NLS-Cre recombinase gene demonstrated recombination in a functional assay *in vitro* (**Fig. S4**). This Cre(intron) recombinase gene was cloned into the LoxP-HBV vectors (**Fig. 2A**) and validated by restriction digest that the full-length plasmid, pCre/LoxP-HBV, was maintained (**Fig. 2B**). The CMV-NLS-Cre cassette utilizes the native HBV polyadenylation signal in the plasmid and upon Cre excision, the polyadenylation signal is absent and replaced with the downstream HDV and hammerhead ribozymes in the 3’ untranslated region, dramatically decreasing Cre expression. After transfection, we verified recombination had occurred by PCR amplifying over the LoxP junction, confirming the correction size by a gel electrophoresis (**Fig. 2C**). The precision of Cre-mediated excision and rcccDNA formation was confirmed by sequencing the amplicon (**Fig. S5**). We next validated that the pCre/LoxP-HBV plasmids were functional *in vivo* using hydrodynamic injections as before. Peak expression levels of HBsAg one week post-injection (range: 4,494 ± 407 ng/mL −6,013 ± 418 ng/mL) were comparable to the two plasmid (*trans*) system, confirming the efficient formation of rcccDNA inside cells with the one plasmid (*cis*) system (**Fig. 2D**).

**Figure 2.**
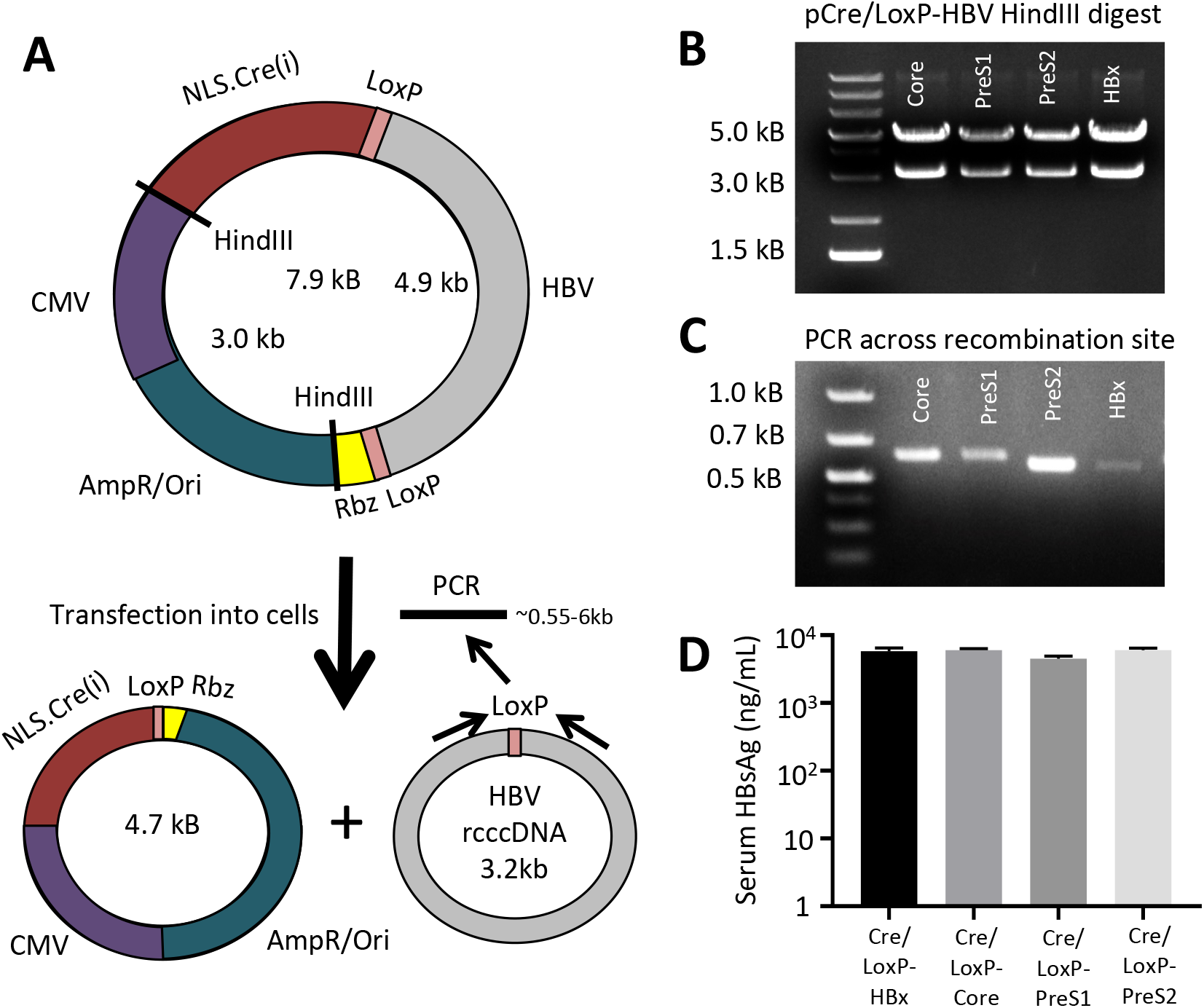
A plasmid with Cre recombinase in cis is stable and forms recombinant cccDNA molecules. **(A)** A CMV.NLS-Cre cassette, with inserted internal intron to inhibit bacterial expression, was cloned into the LoxP-HBV plasmids, such that the Cre transcript would utilize the HBV polyadenylation site. Upon introduction into mammalian cells, expressed Cre will excise the HBV genome forming to separate circular DNA molecules inside the cell. **(B)** The stability of the Cre/LoxP-HBV plasmids in *E. coli* were assessed by HindIII restriction digest, showing that their total size is the 7.9 kB of the original plasmid. **(C)** Upon transfection into 293T cells, recombination can be confirmed using PCR across the recombination junction showing predicted band sizes are present. **(D)** Livers of female NSG mice were transfected with 20 μg Cre/LoxP-HBV plasmids using hydrodynamic tail vein injection and HBsAg levels in the serum assessed week 1 post-injection (n=4). Data represent mean ± standard error of mean (SEM).

### Recombinant cccDNA is regulated by HBx

The biology of rcccDNA was further evaluated by testing the role of HBx protein in its function. HBx is thought to play a crucial in maintaining the open epigenetic state of the cccDNA chromosome [34] and helping to increase transcription of HBV promoters [21]. We asked if HBx is crucial for expression of rcccDNA, which would validate rcccDNA as biologically similar to authentic HBV cccDNA. To this end, we tested the importance of the HBx in activating HBV expression *in vivo* in mice, similar to a previous HBV plasmid study [23]. We utilized the pCre/LoxP-PreS2 construct, since this particular insertion is farthest away from potential HBV gene regulatory elements, and cloned a stop mutation at codon 7 of the HBx ORF (*7, CAA -> TAA) into pCre/LoxP-PreS2, similar to the previous scheme [23]. We found that HBx expression from pSI-X could rescue Core expression in rcccDNA lacking HBx, with no background core expression in pCre/LoxP-PreS2*7 + control (**Fig. 3**). By comparison, pHBV1.2*7 plasmid had rare Core-positive hepatocytes, that increased to wild-type levels with complementation of pSI-X (**Fig. 3**). This indicates that rcccDNA may be similar to authentic infection in absolute requirement for HBx-dependent expression compared to HBV plasmid vectors.

**Figure 3.**
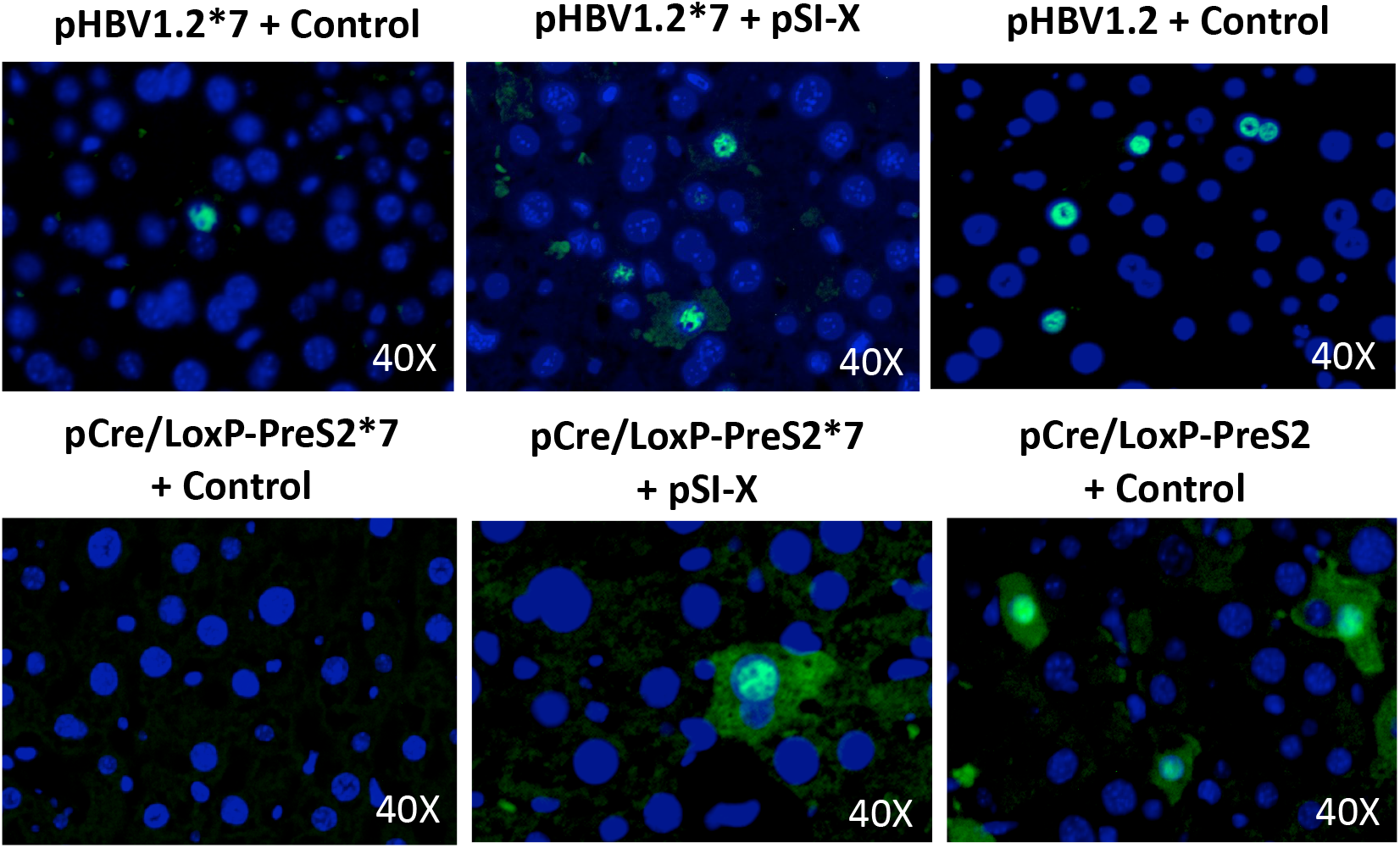
HBx protein regulates HBV core expression in rcccDNA. HBx function was probed by injecting 15 μg pHBV1.2 or pHBV1.2*7 with 5 μg pSI-X expressing HBx or control plasmid (pCMV-Gaussia) into mice. Similarly, 15 μg pCre/LoxP-PreS2 or pCre/LoxP-PreS2*7 with 5 μg pSI-X or control plasmid was injected into mice. At one week post-injection, mice were harvested and tissue analyzed. Immunofluorescent staining for Core protein across the six conditions was undertaken, depicting dependence on HBx for rcccDNA expression. Representative photos from 4 mice are shown.

### Creation of a reporter assay for studying cccDNA levels using a cis-Cre/LoxP plasmid

We next explored using our pCre/LoxP-HBV system to develop a novel drug screen for cccDNA modulating agents. We inserted GFP into the HBV genome over HBsAg, abolishing polymerase expression (pCre/LoxP-HBV-GFP(S); **Fig. 4A**), similar to a previous design [35], which maintains a similar size to wildtype HBV genome. In order to validate the use of this system as a reporter for cccDNA levels, we used the CRISPR-Cas9 in order to degrade cccDNA, utilizing the guide RNA 21 (gRNA21) targeting a conserved site in the HBV genome previously described [26]. Co-transfection of pCre/LoxP-HBV-GFP(S) and pCas9-gRNA21 into 293T cells resulted in decreased GFP levels versus cells transfected with control plasmid (**Fig. 4B**). This serves as a proof of concept for building high-throughput screens for drugs that could trigger cccDNA degradation.

**Figure 4.**
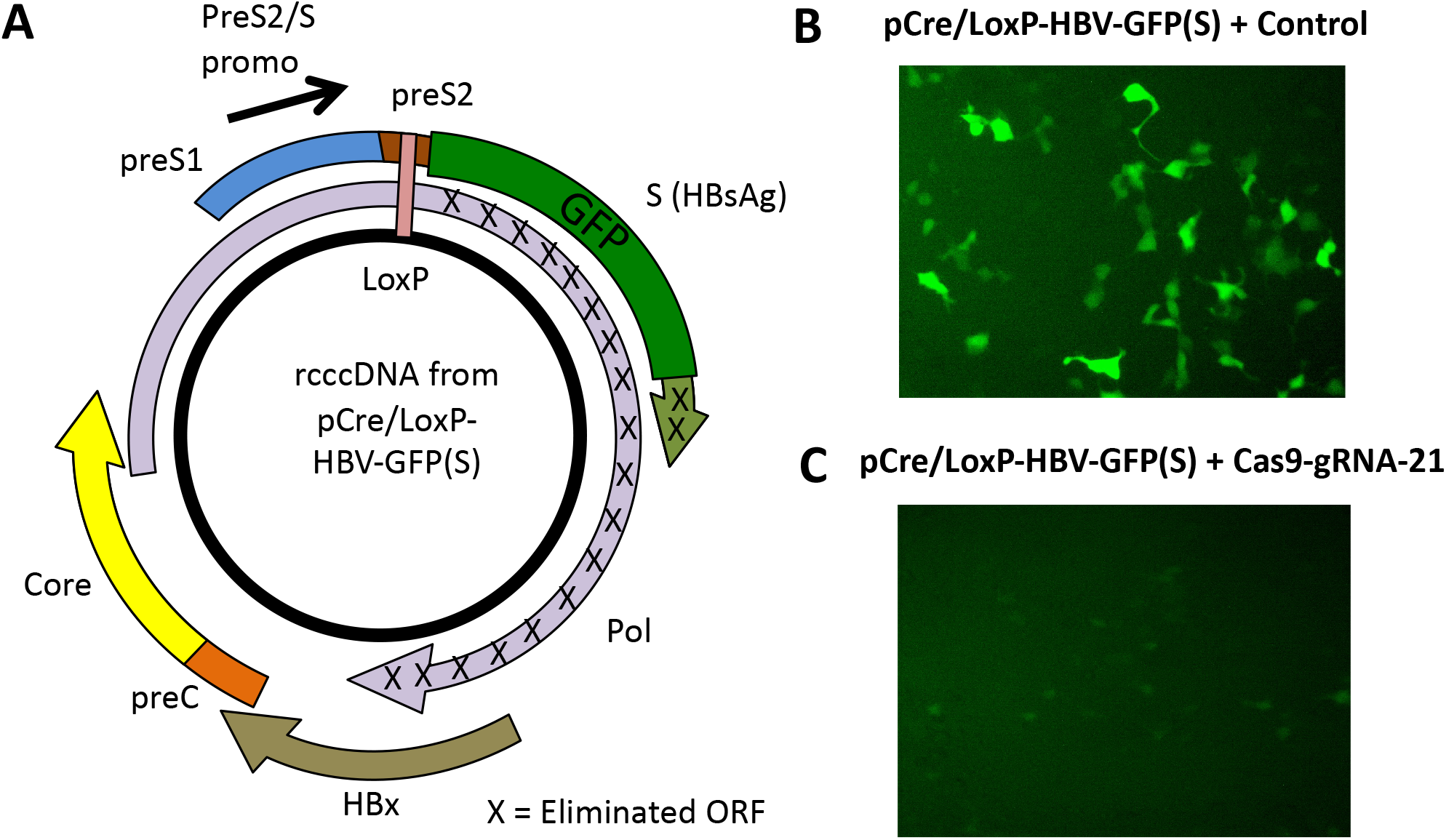
A fluorescent rcccDNA reporter exhibits knockdown after CRISPR targeting. **(A)** Green fluorescent protein (GFP) was cloned into the HBsAg ORF abolishing the overlapping polymerase reading frame. The final molecule after rcccDNA formation is depicted. **(B)** 293T cells were transfected with equal amounts of pCre/LoxP-HBV-GFP(S) and pCas9-gRNA21 or control plasmid. At 48 hours post-transfection, expression of GFP could be detected in 293T cells with control plasmid, whereas transfection with pCas9-gRNA21 revealed greatly reduced GFP expression.

## Discussion

In this study, we described novel research tools for studying cccDNA through artificial recombination methods. We began by identifying four sites in the HBV genome that are tolerant to insertion points without disrupting viral replication. The ability to identify four different sites is beneficial for any studies using rcccDNA vectors, since any one insertion site might influence some feature of HBV function. We were able to define differences between HBV plasmids and rcccDNA in mouse liver over time, demonstrating stable HBsAg expression over 2.5 months, 297-fold higher than pHBV1.3 plasmid at day 78. This difference is crucially important since the HBV plasmid hydrodynamic mouse model has been used widely as a model for testing drugs and cure strategies for HBV [36]. By comparison, rcccDNA was exceptionally stable in our system similar to its behavior in human patients as well [4].

Our results compare favorably to previous artificial recombinase strategies that used different methods and insertion sites in order to generate rcccDNA [7-11]. These rcccDNA molecues also achieved HBsAg levels in the 10,000’s ng/mL after hydrodynamic injection in mice [9]. Moreover, another rcccDNA format showed similar stable HBsAg expression with minimal loss over 49 days after hydrodynamic injection in C3H/HeN mice [8]. These other rcccDNA studies targeted insertions ahead of the precore protein [7], ahead of the PreS1 protein [8], and within an artificial intron inside the HBsAg gene [9]. Given the four additional sites targeted in this study, together there are seven locations now described in the HBV genome that are amenable to manipulation, which will serve as mutual controls for each other.

Innovating on previously published rcccDNA systems, we also developed a novel method of generating rcccDNA with pCre/LoxP-HBV vectors. The plasmids could be propagated stably with both elements, and efficiently formed rcccDNA after transfection. This approach is simpler than using minicircle technologies to generate rcccDNA inside *E. coli*, which require novel bacterial strains and induction media [13]. Furthermore, the cis-Cre recombinase is useful for activating any other floxed reporter genes located in the cell, which can’t be captured with minicircle methods. We utilized this feature in a different study to activate floxed luciferase expression inside rcccDNA-transfected mouse hepatocytes to monitor viability during immune response [37]. For *in vitro* experiments, pCre/LoxP-HBV series generates a homogeneous rcccDNA population, while transfection of pHBV1.3 plasmids results in a mixture of plasmid and cccDNA inside human liver cell lines. Moreover, our pCre/LoxP-HBV plasmids do not require the use of any Cre-expressing mice in order to form rcccDNA [11]. Our novel tools of studying HBx regulation of rcccDNA and the GFP rcccDNA reporter may also find utility in studies of HBV basic biology and drug screening.

In conclusion, this study has established new tools for HBV research, leveraging the artificial generation of cccDNA via Cre recombinase. We describe the generation of plasmids that can convert into recombinant HBV cccDNA upon transfection. Ultimately, we believe the versatility of these tools will benefit the field in more easily establishing chronic HBV infection models and studying questions of viral function and immune clearance.

## Acknowledgements

RLK was supported by NIH grant T32DK060445. Thanks go to Urtzi Garaigorta and Christina Whitten-Bauer for assistance with the HBV qPCR and ELISA assays, Betty Slagle for providing the HBx plasmids, and Karl-Dimiter Bissig for advice during the project.

## Declaration of competing interest

The authors disclose no potential conflicts of interest.

## Notes

### Competing Interest Statement

The authors have declared no competing interest.

### Summary of Updates

Figure 1 and 2 revised.

