## Supplemental Figures for "Cre/LoxP-HBV plasmids generating recombinant covalently closed circular DNA genome upon transfection"

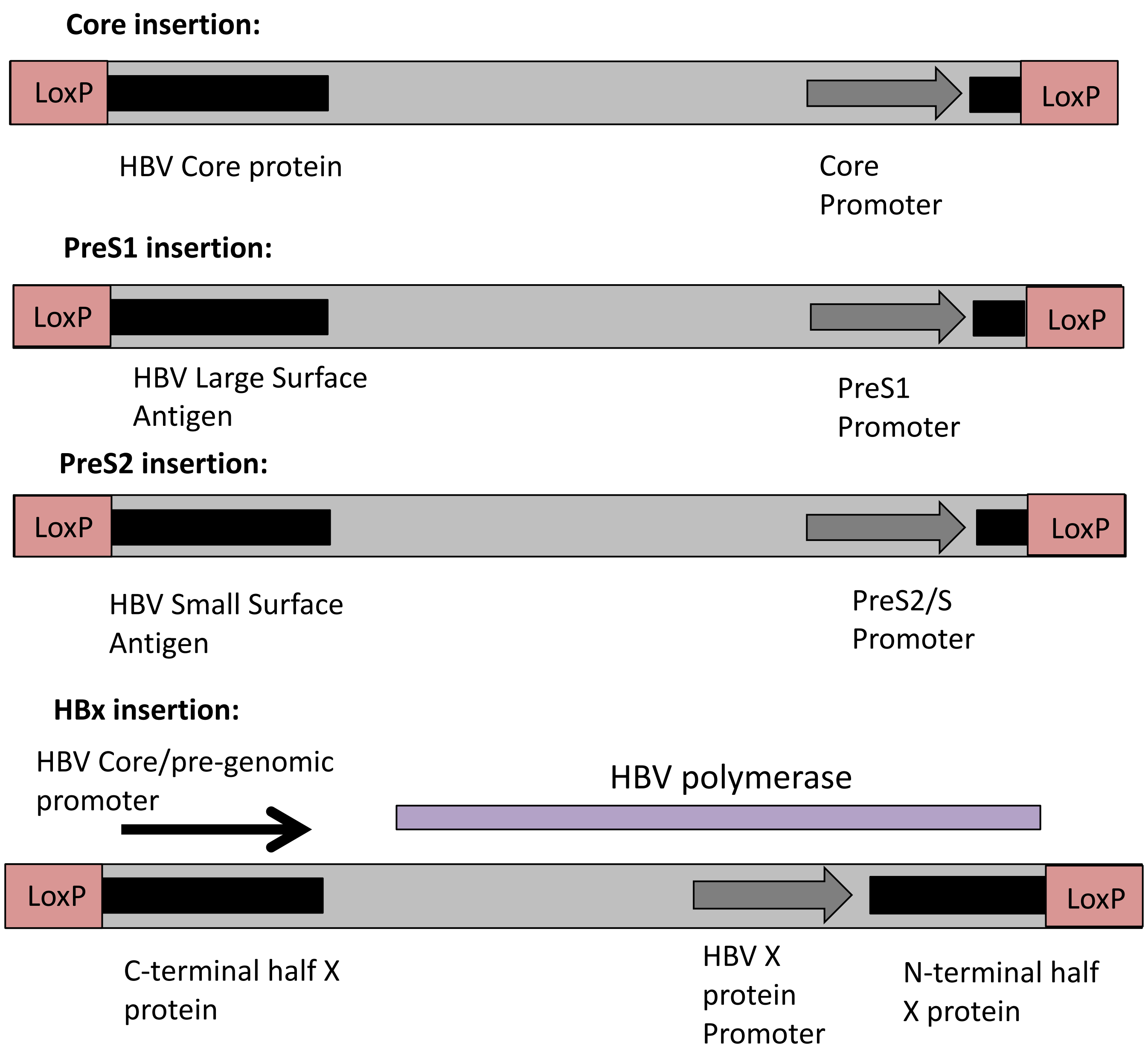

**Supplemental Figure 1. Schematics and sequence information for LoxP-HBV vectors.** Schematics representing the four different insertion points in their context in the linear LoxP-HBV vector are depicted. For each, the HBV protein is interrupted when in the linear context, leaving only a small n-terminal half still expressed. For the LoxP-HBx cassette, it is noted that the HBV polymerase ORF ends right before the LoxP insertion, and that the HBc promoter is located after the LoxP site in order to drive expression. Black = viral protein ORF ; dark grey = promoter ; purple = HBV polymerase protein.

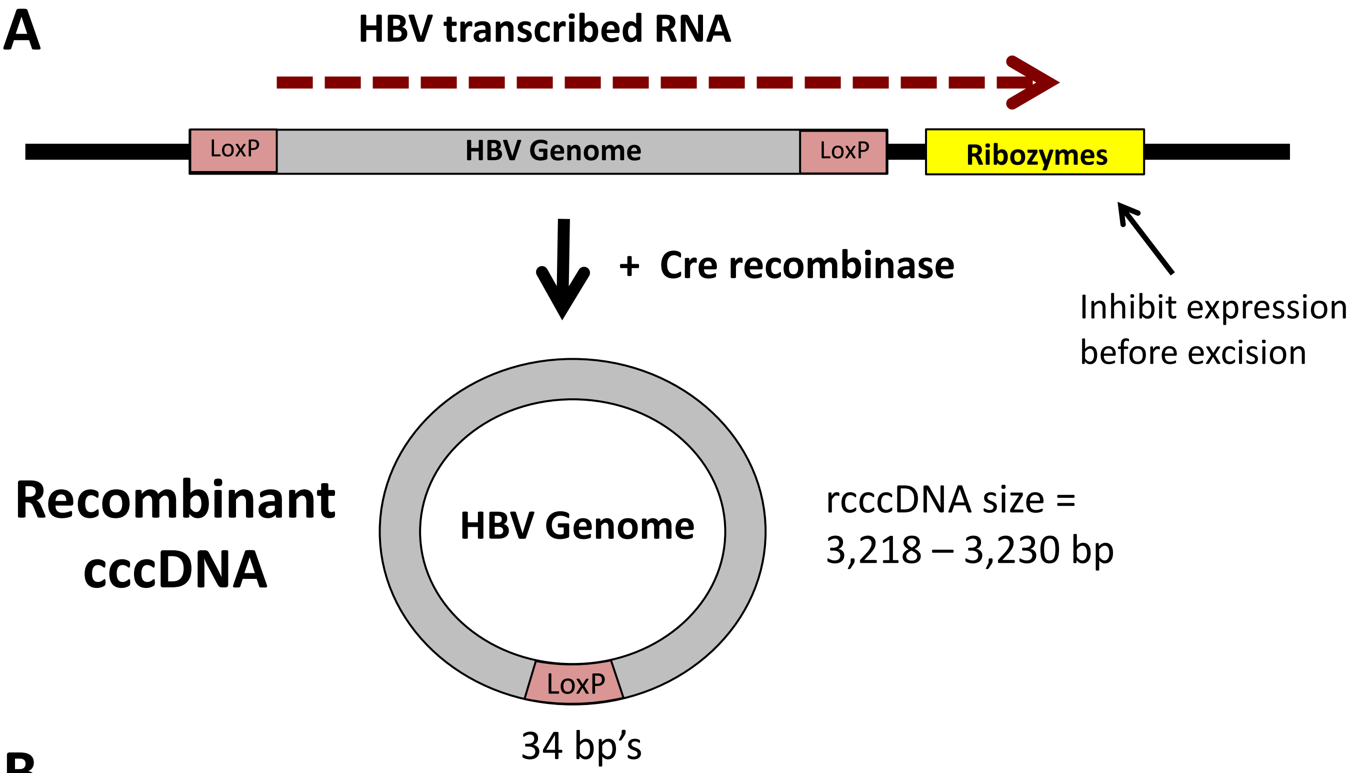

**B**

CCgcGG-AAACAAACAAA-
*CTGAGATGCAGGTACATCCAGCTGATGAGTCCCAAATAGGACGAAACGCGCTTCGGTG*
*CGTCCTGGATTCCACTGCTATCCAC* -AAAAAGAAAATAAAACAAA -
CAGATGGCCGGCATGGTCCCAGCCTCCTCGCTGGCGCCGGCTGGGCAACATTCCGAGG
GGACCGTCCCCTCGGTAATGGCGAATGGGACCC – AAAGAAAGAAA – GCTAGC

**C**

**Core:** M D **I T S Y S I H Y T K L S** I D P Y K E F G A T V E L

**PreS1:** M G G W S S **T G I T S Y I L S Y T K L S G T** F R A N T

**PreS2:** M Q W N S T **I T S Y I L S Y T K L S** L Q D P R V R G L

**HBx:** R M E T T V **I T S I T S Y S I H Y T K L S** N A H Q I L P

**Supplemental Figure 2. Cre recombinase mediates excision to form recombinant cccDNA and sequence information for the pLoxP-HBV plasmids.**
(**A**) The LoxP-flanked HBV genomes were cloned into a plasmid vector with downstream self-cleaving ribozyme sequences in order to promote transcript degradation and inhibit protein expression. Upon introduction of Cre recombinase, the LoxP-HBV cassette will be excised from the plasmid vector and form a recombinant HBV cccDNA molecule. (**B**) The ribozyme sequence located 3' to the LoxP-HBV cassette to disrupt HBV expression is provided. The sequence consists of a SacII restriction site – flexible RNA linker - *Schistosoma mansoni* hammerhead ribozyme (italics) – flexible RNA linker – hepatitis delta virus ribozyme (underlined) – flexible RNA linker – NheI restriction site. (**C**) The amino acid sequences for the LoxP sites inserted into the open reading frames of the designated HBV proteins are provided in red; in green, amino acid sequence of the introduced restriction sites for the PreS1 construct.

**LoxP - Core**

AAG CTG TGC CTT GGG TGG CTT TGG GGC **ATG** GAC ATA ACT TCG TAT AGC ATA CAT TAT

ACG AAG TTA TCC ATC GAC CCT TAT AAA GAA TTT GGA GCT ACT GTG GAG TTA CTC TCG TTT

**LoxP – PreS1**

CAC CAT ATT CTT GGG AAC AAG ATC TAC AGC **ATG** GGA GGT TGG TCA TCG **ACC GGT** ATA ACT TCG TAT ATT CTA TCT TAT ACG AAG TTA TCT **GGT ACC** TTC AGA GCA AAC ACC GCA AAT CCA GAT TGG GAC TTC AAT CCC AAC AAG GAC ACC

**LoxP – PreS2**

CAG CCT ACC CCG CTG TCT CCA CCT TTG AGA AAC ACT CAT CCT CAG GCC **ATG** CAG TGG AAT TCC ACA ATA ACT TCG TAT ATT CTA TCT TAT ACG AAG TTA TCT CTG CAA GAT CCC AGA GTG AGA GGC CTG TAT TTC CCT GCT GGT GGC TCC

**LoxP – HBx**

CCC GTC TGT GCC TTC TCA TCT GCC GGA CCG TGT GCA CTT CGC **TTC ACC TCT GC**A CGT CGC ATG GAG ACC ACC GTG ATA ACT TCG TAT AGC ATA CAT TAT ACG AAG TTA TCC AAC GCC CAC CAA ATA TTG CCC AAG GTC TTA CAT AAG AGG ACT CTT GGA CTC TCA GCA

**Supplemental Figure 3. Complete DNA sequence information for LoxP insertions into the different HBV genomic sites.**
DNA sequence information is provided for the Core, PreS1, PreS2, and HBx insertion sites in the HBV genome, with sequence representing the recombined molecule with HBV sequence on both sides of the LoxP site. Note that the PreS1 and PreS2 insertions contain an alternative LoxP sequence from the wildtype in order to maintain both surface antigen and polymerase open reading frames. Underlined = LoxP site, Red = Start codon for HBV protein; Green = additional inserted restriction sites (5’ AgeI, 3’ KpnI); Brown = DR2 sequence. Purple = Stop codon for HBV polymerase

**A**

**ACGTGCAAAACAG***gtaagcgcccctaaaatccctttggcacaatgtgtcctgaggggA*
*gaggcagcgacctgtagatgggacgggggcactaaccctcagggtttggggttctgaatg*
*tgagtatcgccatgtaagcccagtatttggccaatctcagaaagctcctggctccctggagg*
*atggagagagaaaaacaaacagctcctggagcagggagatgctggcctcttgctctccgg*
*ctccctctgttgccctctggtttctccccag***GCTCTAGCGTTCGAACGCACTGATTTC**
**GA**

**B**

Insertion point in the Cre amino acid sequence: $^{130}$RAKQ – *intron* – ALAF$^{137}$

**C**

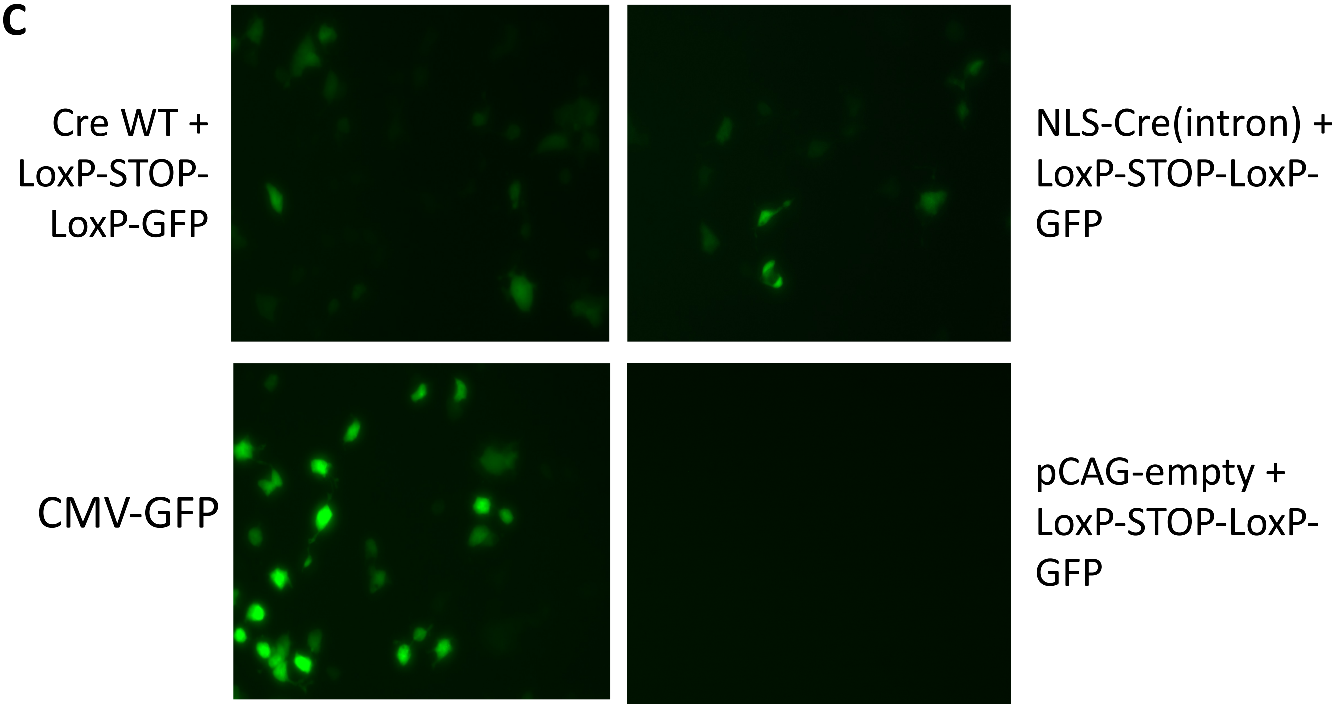

**Supplemental Figure 4. Cre recombinase gene with an inserted mammalian intron into the coding sequence was designed and shown to be functional in mediating recombination.** (**A**) The DNA sequence for Cre recombinase (bold, capitalized) with the insert intron from the human growth hormone (italics, lower case) is provided. The site was selected based on the canonical CAG/G recognition sequence for mammalian introns. (**B**) The amino acid sequence around the intron insertion point between amino acid 133Q and 134A in the Cre recombinase open reading frame) is provided. (**C**) In order to validate the intron was processed and functional Cre recombinase was made, we co-transfected a pLoxP-STOP-LoxP GFP reporter vector with pCMV-Cre(intron), pCMV-Cre, or control plasmid in equal amounts into 293T cells. A transfected pCMV-GFP vector served as a positive control. Fluorescent imaging for GFP expression was taken 48 hours post-transfection.

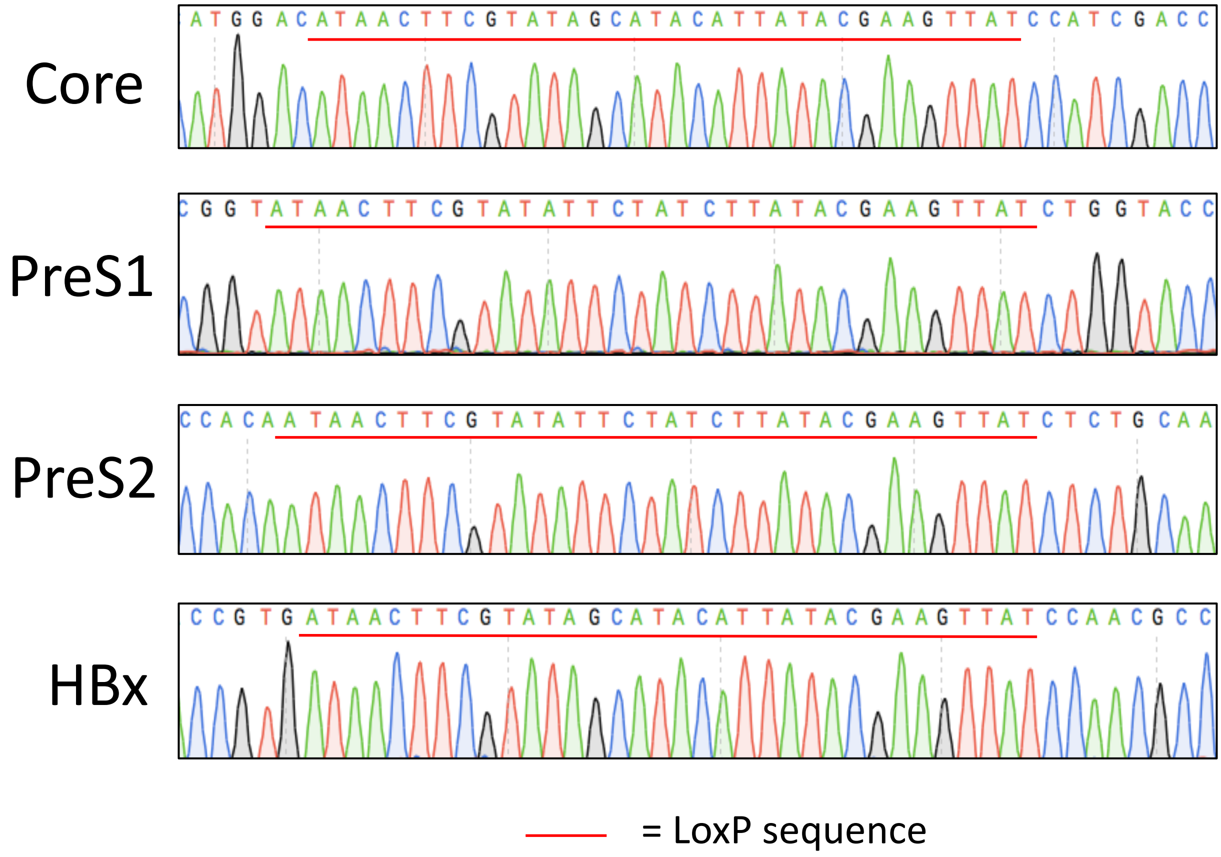

**Supplemental Figure 5. Sequencing of LoxP junction confirms proper recombination of Cre/LoxP-HBV plasmids.**
The pCre/LoxP-HBV plasmids were transfected into 293T cells allowing recombination to occur intracellularly. A PCR reaction using primers that amplify the LoxP junction after Cre-mediated recombination was performed. The bands were sequenced for the different LoxP insertions confirming formation of rcccDNA with the expected HBV sequence on either side. The sequence of the LoxP site is underlined in red.

| Primers used to amplify over cccDNA recombination junction in 293T cells | |
|---|---|
| LoxP-HBx For | TAGGCTGTGCTGCCAACTGGATC |
| LoxP-HBx Rev | GATGTCCATGCCCCAAAGCCAC |
| LoxP-Core For | ACTTCGCTTCACCTCTGCACGTC |
| LoxP-Core Rev | TAGGTCTCTAGACGCTGGATCTTCC |
| LoxP-PreS1 For | GTTTGTAGGCCCACTCACAGTTAATGAG |
| LoxP-PreS1 Rev | GATGAGTGTTTCTCAAAGGTGGAGACAG |
| LoxP-PreS2 For | GAGCAAACACCGCAAATCCAGATTGG |
| LoxP-PreS2 Rev | CTGCGAATTTTGGCCAAGACACACG |
| **Primer used to make the LoxP-HBV plasmids** | |
| PreS1 insert Rev 1 | TTTCCGCGGAGATAACTTCGTATAAGATAGAATATACGAAGTTATACCGGTCGATGACCAACCTCCCA |
| PreS1 insert For 2 | AAACCCGGGATAACTTCGTATATTCTATCTTATACGAAGTTATCTGGTACCTTCAGAGCAAACACCGCA |
| HBx insert For 1 | AAACCCGGGATAACTTCGTATAGCATACATTATACGAAGTTATCCAACGCCCACCAAATATTGCCCAAGG |
| Core insert For 1 | AAACCCGGGATAACTTCGTATAGCATACATTATACGAAGTTATCCATCGACCCTTATAAAGAATTTGGAGCTACTG |
| HBx/Core insert Rev 1 | TCCCCAATCCTCGAGAAGATTGACG |
| HBX/Core insert For 2 | CGTCAATCTTCTCGAGGATTGGGGA |
| Core insert Rev 2 | TTTCCGCGGATAACTTCGTATAATGTATGCTATACGAAGTTATGTCCATGCCCCAAAGCCACCCAA |
| S1/S2 insert For 1 | CGTTTCCATGGCTGCTAGGCTG |
| PreS2 insert Rev 1 | TTTCCGCGGAGATAACTTCGTATAAGATAGAATATACGAAGTTATTGTGGAATTCCACTGCATGGCCTG |
| PreS2 insert For 2 | AAACCCGGGATAACTTCGTATATTCTATCTTATACGAAGTTATCTCTGCAAGATCCCAGAGTGAGAGG |
| S1/S2 insert Rev 2 | CAGCCTAGCAGCCATGGAAACG |
| HBx insert Rev 2 | TTTCCGCGGGGATAACTTCGTATAATGTATGCTATACGAAGTTATCACGGTGGTCTCCATGCGAC |
| **Construction of the Cre/LoxP-HBV/GFP vectors** | |
| GFP into SAg For | ACAGTAGAATTCCATGGTGAGCAAGGGCGAGGAG |
| GFP into SAg Rev | GATGACACTAGTTACTTGTACAGCTCGTCCATGCCGA |

**Supplemental Table 1. Primers utilized in this study are provided.**
Primers for generating the LoxP-HBV plasmids starting with a pHBV1.3 ayw strain genotype D template are given. Primers utilized to amplify over the LoxP junction to form cccDNA are provided for both *in vitro* and *in vivo* experiments. Primers for generation of pCre/LoxP-HBV-GFP(S) are also provided.
